# How Our Cells Become Our Selves: The Cellular Phylodynamic Biology of Growth and Development

**DOI:** 10.1101/2021.09.22.461268

**Authors:** Philip Chodrow, Jessica Su, Daniel Lee, Tahmid Ahmed, Neil He, De Man Ruben, Ankur Tiwari, William Mannherz, Luca Citi, Daniel DiCorpo, James Selib Michaelson

## Abstract

Our lives begin with 1 cell, then 2, then 4, then the trillion cell adult, comprised of cell lineages, tissues, organs. How does this occur? Examination in numbers of cells, ***N, Cellular Phylodynamics***, revealed two previously unappreciated processes: ***UNI-GROWTH***, the slowing of growth that occurs as we become larger, caused by fewer cells dividing, captured by the ***Universal Mitotic Fraction*** and ***Universal Growth Equations***, with accuracy confirmed for 13 species, including nematodes, mollusks, and vertebrates; and ***ALLO-GROWTH***, the creation of body parts from ***Founder Cells***, captured by the ***Cellular Allometric Growth Equation***, which describes mitotic expansion by ***Cell-Heritable*** change in the ***Cell Cycle Time***. These equations can generate cell lineage approximations, bringing the power of coalescent theory to developmental biology.

## Introduction

Our lives begin with 1 fertilized cell, dividing to become 2 cells, then 4, then 8, growing to the trillion-or-so-celled human^1^ or the thousand-or-so-celled nematode^2^, comprised of cell lineages, tissues, organs, and anatomical structures, of recognizable size and shape. How does this occur? When considered in aggregate, this appears to us as growth, a process with practical implications in agriculture, aquaculture, and health.^3–6^ When considered in detail, this appears to us as embryonic development, whose mechanisms await our decipherment.

Examination of animal growth and development in continuous units of length/volume/mass has yielded quantitative insights, whose meanings have often been unclear. For example, while growth starts out exponentially, it is soon tempered by a decline in rate, and many ***density dependent growth equations***, which form ***S-shaped*** growth curves, have been considered^1–9^, none of which accurately fit the growth of real animals.^10–12^ Within the body, the relative growth of tissues, organs, and anatomical structures often appears as straight lines on log-log graphs, although the cause of such ***allometric growth*** has long been a mystery.^13,14^ ***Cellular-Selection***, that is, differential cellular proliferation and death, creates many anatomical structures by outgrowth from the undifferentiated body mass, and molds much of the fine scale detail of anatomy, although how cell division is harnessed to achieve this is obscure.^15,16^ Cell lineages grow from individual ***Founder Cells***,^2,17^ although how embryos use mitosis to create these parts,^18,19^ and how these ***Founder Cells*** undergo ***Cell-Heritable*** change that characterized each clone,^16,35^ remains unknown. The analysis of cellular diversification, by single cell mRNA expression,^50,53,54^ cell marking,^21,20,51^ and high resolution 4D microscopy,^18,19,47^ has also wrestled with the problem of cell lineage formation.^21^

Here, we report the examination of growth and development in discrete units of numbers of cells, ***N***. We call this approach “***Cellular Phylodynamics***”. Our frame of mind is inspired by Stadler, Pybus, and Stumpf,^22^ who have articulated how ***Phylodynamics***, a powerful approach widely used in epidemiology, taxonomy, and population biology ^23–30^ can also be brought to bear in the study of developmental biology. As they stated, “The term ‘phylodynamics’ describes the integration of phylogenetic and population dynamic models to study tree-generating processes”, and that such “‘***tree-thinking***’ … can be beneficial to the interpretation of empirical data pertaining to the individual cells of multicellular organisms.” As we shall see, the application of such a ***Cellular Phylodynamic tree-thinking*** approach to the analysis of data on numbers of cells in animals as a whole, ***Nw***, and in their parts, ***N***_***p***_, does indeed provide new insight into how we grow and develop.

## Results

### UNI-GROWTH

#### The problem of body growth and development

For more than two centuries, biologists have appreciated that growth begins exponentially, but then declines in rate, and many expressions to capture such ***density dependent growth*** have been considered,^3–12^ none of which accurately captures the growth of real animals.^11,12^ Our goal here is not to search for another equation that fits the growth of animals, but to carry out a ***Cellular Phylodynamic Analysis*** to examine what actual growth data, in units of numbers of cells, ***N***, can tell us about the nature of the growth and development of the body.

#### The ***Cellular Phylodynamic Analysis*** of body growth and development

To carry out ***Cellular Phylodynamic Analysis*** of the growth of the body as a whole, we assembled values on the numbers of cells, ***N***_***w***_, from fertilization until maturity, for nematodes, frogs, chickens, cows, geese, quail, turkeys, zebrafish, seabass, clams, mice, rats, humans, and other animals (FIGURE 1[left], and FIGURES A2, A3, and A5 in APPENDIX). A detailed account of how these values were assembled can be found in the APPENDIX.

**FIGURE 1:**
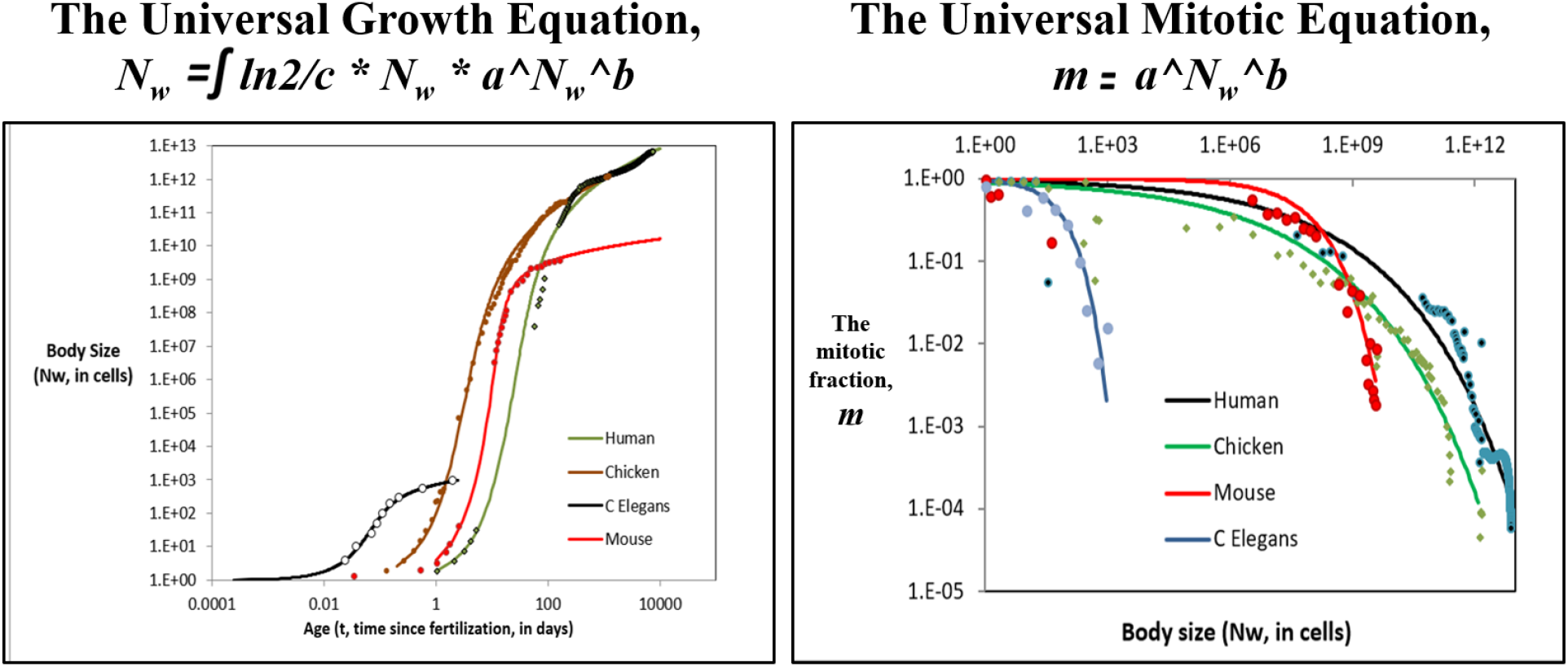
Growth data and the *Universal Growth Equation* and *Universal Mitotic Fraction Equation for capturing growth from fertilization until maturity*. **Left: Growth, in units of numbers of cells, *N*_*w*_, from fertilization until maturity, in units of days, *t*.** Datapoints for animal size’s (***N***_***x***_) versus time (***t***), for ***C elegans*** chickens, mice, turkeys, quail, geese, frogs, and humans, and their fit to the numerically integrated form of the ***Universal Growth*** ***Equation*** (#5b). For graphs of the growth of other species, see FIGURE A1, A2, A5, and A6 in APPENDIX. **>Right: Estimate of the decline in the *Mitotic Fraction*, a measure of the fraction of cells dividing that occurs as animals increase in size, as estimated by the *Mitotic Fraction Method* and captured by the *Universal Mitotic Fraction Equation***. Data points for animal size, ***N***_***w***_, in integer units of numbers of cells, vs. the ***Mitotic Fraction, m***, from fertilization, until maturity, for humans, chickens, ***C elegans*** nematode worms, and mice. For additional graphs of the change in the ***Mitotic Fraction, m***, with size, ***N***_***w***_, see FIGURE A2, A3, and A5 in APPENDIX.

Many studies have noted that mitosis occurs in the great majority of cells at the beginning of development, giving early embryos the capacity to increase in size, ***N***_***w***_, with age, ***t***, such that the ***rate of growth*** is:

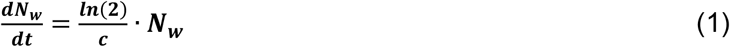

Where ***N***_***w***_ is the number of cells in the body as a whole, and ***c*** is the ***Cell Cycle Time***. Integration shows that when all cells are dividing, growth is exponential:

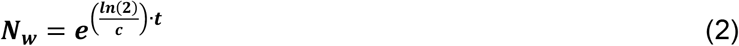

Such exponential growth can be visualized by graphing the log of the numbers of cells, ***N***, against time, that is:

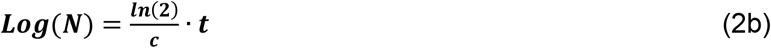

Thus, data on the relationship between the log of cell number, ***Log***(***N***_***w***_), and time, ***t***, can be characterized at the very beginning of development, when all cells are dividing, to give one a practical measure of the ***Cell Cycle Time, c***. (See APPENDIX for details). Values for ***c*** for each of the 13 animals noted above can be seen in the TABLE.

This early unrestricted growth soon gives way to a decrease in the fraction of cells dividing, (for references, see APPENDIX) driven by ***mitotic quiescence***, the complex biochemical mechanism within cells that prevents mitosis from occurring in response to inhibitory and stimulatory signaling molecules that cells use to communicate.^31–33^ This decrease in mitotic activity that occurs as we become larger, which we call the “***Mitotic Fraction***”, ***m***, can be taken into account by inserting it into Equation #1:

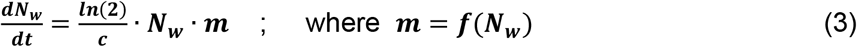

Thus, converting the ***exponential growth*** ***equation*** (#1) into a ***density dependent growth*** ***equation*** (#3). This “***Swiss Army Knife***” of an expression (since ***f***(***N***_***w***_) can accommodate any change in the rate of growth that might occur as an animal becomes larger) balances the unlimited quality of exponential growth that occurs at the beginning of development with the inevitable reduction in growth rate that occurs as we grow.^5^ The result is a classic ***S-shaped*** growth curve, such as can be seen with the data we have assembled on growth of the 13 animals listed above (FIGURE 1, and FIGURES A2, A3, and A5 in APPENDIX).

#### The *Mitotic Fraction Method*

Rearranging Equation #3 allows us to use growth data to characterize how the value of ***Mitotic Fraction, m***, changes as animals increase in size, ***N***_***w***_:

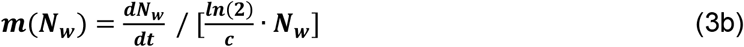

We call this approach the “***Mitotic Fraction Method***”. In simple terms, Equation #3b gives us a rough measure how many cells would have to divide to account for the amount of growth that occurs over each period of time. Details for the practical execution of the method, and the limits of its approximation, can be found in the APPENDIX.

#### The *Universal Mitotic Fraction Equation*

We applied the ***Mitotic Fraction Method*** (Equation #3b) to the growth data for all of the 13 animals listed above. This revealed exponential growth during the first few mitoses (***m*≈1**), smoothly transitioning to an accelerating decrease in the fraction of cells dividing as adulthood approaches (***m*<1**) (FIGURE 1[right], FIGURES A4-A6 in APPENDIX). The basis for the relationship between ***m*** and ***N***_***w***_ was revealed by graphing **ln**(***N***_***w***_) vs. **-ln(-ln**(***m***)), which yielded straight rows of datapoints (FIGURE A5 in APPENDIX), that is:

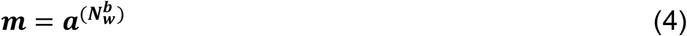

This expression was found to capture, with high ***r***^2^ values, the decrease in the value of the ***Mitotic Fraction, m***, that occurs as animals increase in size, ***N***_***w***_, from fertilization until maturity, for all 13 of species noted above (TABLE). For this reason, we call it the “***Universal Mitotic Fraction*** ***Equation***” (#4).

#### The *Universal Growth Equation*

Combining Equations #3 and #4 leads to this expression for describing the rate of animal growth:

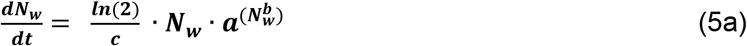

The relationship between size, ***N***_***w***_, and age, ***t***, is captured by numerical integration of Equation #5, where the age, ***t***, at fertilization is ∼0, and the age at death is ***d***:

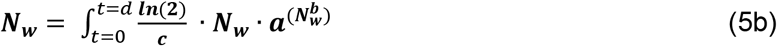

as well as by this closed-form integration:

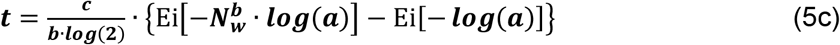

where **Ei** is the exponential integral function.^34^

We tested the numerically integrated form of these expressions, Equation #5b, on body size, ***N***_***w***_, vs age, ***t***, from fertilization until maturity, for all 13 species listed above, finding close fits, i.e., high ***r***^2^ values (FIGURE 1[left], TABLE). For this reason, we call these expressions “***Universal Growth Equations***”. Tests of other widely used density dependent growth equations found none to fit actual growth data as well as the ***Universal Growth Equation*** (FIGURES A3-A5 in APPENDIX). Thus, the ***Universal Growth Equation*** is not only more accurate than previous growth equations, it also succeeds, as previous equations did not, in capturing growth over its full span, from the first fertilized cell of the embryo up to mature size. We call the biological process captured by the ***Universal Mitotic Fraction*** and ***Universal Growth Equations*** “***UNI-GROWTH***”.

#### Possible mechanisms

The ***a*** and ***b*** parameters of the ***Universal Growth Equation*** determine the “curviness” of the “***S*”** of the ***S-shaped*** growth curve, while its parameter ***c***, the ***Cell Cycle Time***, expands or contracts the “***S*”** like an accordion. The ***Universal Mitotic Fraction Equation***, acting within that expression, describes the decline in the ***Mitotic Fraction, m***, that occurs as we grow, as made specific for each animal by the values of its ***a*** and ***b*** parameters. Fortunately, the forms of these expressions give us some hints of how the biochemistry of the cell may make such ***UNI-GROWTH*** possible.

#### The *Mitotic Fraction* and mitotic signaling

What might be the mechanism behind the change in the ***Mitotic Fraction, m***, that occurs as we grow? A tempting possibility emerges from the mathematical analysis of the deceptively simple discrete allocation of ligand molecules among cells (Equations #700-#720, APPENDIX).^35,36^ We examined the consequences for cells growing in a confined space, such as an egg or uterus, if these cells produce inhibitory growth factor molecules. As the number of cells increase, the concentration of the inhibitory growth factor increases as well, and the fraction of cells that have bound one or more of these factors also increases. Most remarkably, the math reveals that the decline in the fraction of cells that had bound one or more of these inhibitory factors occurs by an expression of exactly the same form as the ***Universal Mitotic Fraction Equation*** (#4). Thus, little more than the binding of growth factor molecules to cells, driven by conventional thermodynamics, is sufficient to account for the decline in the fraction of cells dividing that occurs as we grow, a hypothesis-generating possibility, worthy of experimental analysis.

#### *Cell Cycle Time, c*, growth, junk DNA

The ***Cell Cycle Time, c***, of the 13 animals we have examined here ranges from just 15 minutes for nematodes to about a day for humans (TABLE). Genome size is known to be the main determinant of ***Cell Cycle Time***, as first reported by Van’t Hof and Sparrow,^37^ for plant cells, subsequently confirmed for cells from many types of organisms (see APPENDIX for references and details). Apparently, the more DNA a cell has to copy, the longer it takes to divide. Animal genomes are usually comprised principally non-coding DNA,^38^ mostly transposable elements that are very susceptible to duplication and deletion,^39–43^ and thus capable of providing abundant genetic variation in genome size. This genetic variation in genome size should offer genetic variation in ***Cell Cycle Time***, and thus variation in growth rate, which the bitter reality of Darwinian selection could draw upon to allow species to drift to growth rates that give them their greatest chance of survival. Could our much maligned “junk DNA” owe its existence to this important function?

### ALLO-GROWTH

#### The problem of body part growth and development

For more than a century, biologists examining the growth of various parts of the body, ***p***, in comparison to the size of the body as a whole, ***w***, in continuous units of weight, volume, and length, have frequently found straight lines on log-log graphs. This observation is called ***allometric growth:***^13,44,45^

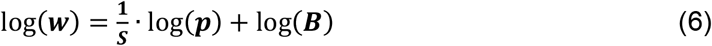

The reasons why the growth of the parts of the body should take this form, and the meaning of the parameters of this ***Allometric Growth Equation***, have long been mysteries.^44,45^ Our goal here is to carry out a ***Cellular Phylodynamic Analysis*** to examine what actual growth data, in units of numbers of cells, ***N***, can tell us about the nature of the growth and development of the parts of the body.

#### The ***Cellular Phylodynamic Analysis*** of body part growth and development

To carry out ***Cellular Phylodynamic Analysis*** of the growth and development of the cell lineages, tissues, organs, and anatomical structures of the body, we assembled values, from the literature, on the numbers of cells in the body, ***N***_***w***_, and in its various parts, ***N***_***p***_. A detailed account of how these values were assembled can be found in the APPENDIX.

The anatomical structures of developing nematodes (***Caenorhabditis elegans***^2^ and ***Meloidogyne incognita***^17^) and tunicates (***Oikopleura dioica***^46^), who are chordate quite closely related to ourselves, have been characterized for every cell from the first single cell zygote onward. From these cell lineage charts, we assembled values of ***N***_***p***_ and ***N***_***w***_ (FIGURES 2 and A8-A11 in the APPENDIX).

**FIGURE 2:**
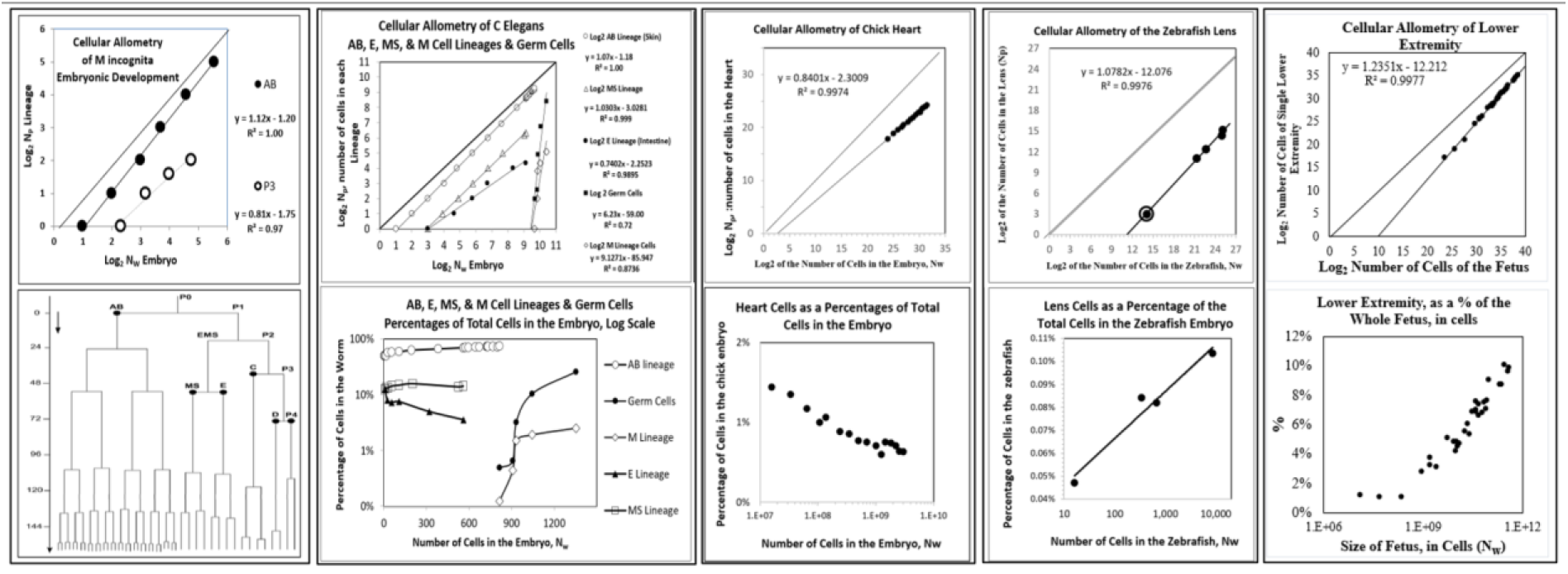
*Cellular Phylodynamic Analysis* of cell lineages, tissues organs, and anatomical structures, and their fit to the *Cellular Allometric Growth Equation*. ***Cell lineages*: *M incognita*** (**AB** and **P** lineages and its cell lineage chart from which these values were derived). ***C elegans*** (**AB, M, E, MS** and **Germ Cell** lineages). The **AB** lineage of nematodes forms the skin while the **E** lineage forms the intestine. ***Tissues, organs***, and ***anatomical stuctures*** of developing fetuses: Chick (heart), Zebrafish (lens), Human (brain), Human (leg)). For references and additional values for growth of the ***cell lineages*** of the nematodes, ***M incognita*** and ***C elegans***, and the chordate tunicate ***O dioica***, and for the ***tissues, organs***, and ***anatomical stuctures*** of the developing fetuses see FIGURES A9-17 in APPENDIX.

The tissues, organs, and anatomical structures seen later in development have usually been measured in units of weight or volume. From these batch values, we could generate useful approximations of ***N***_***p***_ and ***N***_***w***_ (see APPENDIX). We assembled values for chicks (gizzards, livers, hearts, kidneys), rats (livers, brains, kidneys, forelegs, ears, stomachs, spinal cords), humans (brains, livers, kidneys, lungs, pancreases, adrenals, thymuses, spleens, lower extremities, upper extremities, stomach, hearts, intestines), zebrafish (eye lens), mice (livers, brains, kidneys, forelegs), clams, and goldfish (FIGURES 2 and A12-A17 in the APPENDIX).

#### The *Cellular Allometric Growth Equation*

When we placed, on log-log graphs, the values we assembled on the number of cells in cell lineages, tissues, organs, and anatomical structures, of the body, ***N***_***p***_ versus the number of cells in the body as a whole, ***N***_***w***_, in the great majority of cases, these datapoints appeared as straight rows of dots (FIGURE 2 and FIGURES A8-A17 in APPENDIX), as attested by high ***r***^***2***^ values, or:

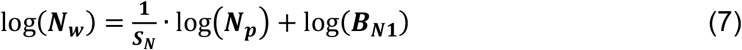

We call this expression the “***Cellular Allometric Growth Equation***” (FIGURE 2), and the biological process that it captures “***ALLO-GROWTH***”. We call the parameter ***B***_***N***_ the “***Cellular Allometric Birth***”’ and ***S***_***N***_, the “***Cellular Allometric Slope***”.

#### Cells to Structures: The *Cellular Allometric Birth, B_N_*

The value of ***B***_***N***_, the ***Cellular Allometric Birth***, can be seen easily on log-log graphs, as the place where the ***Cellular Allometric Growth Equation*** crosses the ***x***-axis. Thus ***B***_***N***_ corresponds to the value of ***N***_***w***_ where **log(*N***_***p***_**)=0**, which is where ***N***_***p***_**=1** (FIGURE 3).

**FIGURE 3:**
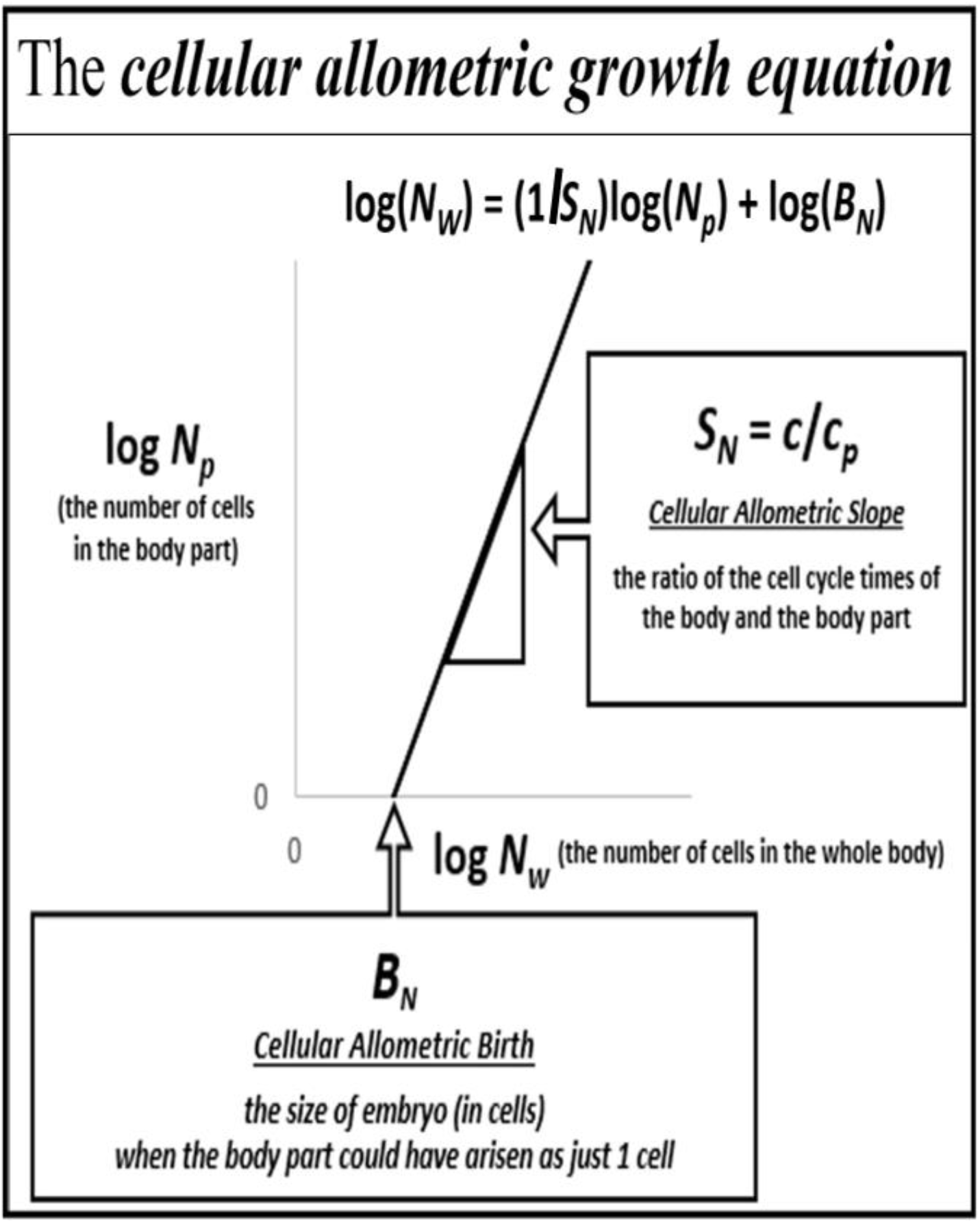
The *Cellular Allometric Growth Equation*.

Should a part of the body come into existence as a single ***Founder Cell***, the ***Cellular Allometric Birth, B***_***N***_, corresponds to the number of cells in the body of the embryo, ***N***_***w***_, when that single ***Founder Cell*** arose by mitosis (i.e., ***B*_*N*_** = ***N*_*w*_** when ***N*_*p*_** = **1**). In fact, this is precisely what has been seen for those animals for which we have ***cell lineage charts***, which have been collected to describe every cell in the embryo from the zygote onward: ***Caenorhabditis elegans***^2^ and ***Meloidogyne incognita***^17^ nematode worms, and ***Oikopleura dioica*** tunicates,^46^ which are chordates quite closely related to ourselves (FIGURE 2 and FIGURES A9-A11 in APPENDIX).

For the tissues, organs, and anatomical structures for which we have growth data later in development, we do not have data on their cellular origins. There has been much scholarship on this point (reviewed in the APPENDIX), but, fortunately, modern light sheet 4D microscopy should allow us to answer these questions,^18,19,47^ and the ***Cellular Allometric Growth Equation*** points us to where we should look. Furthermore, whether a body part arises from a single ***Founder Cell***, or multiple ***Founder Cells, Cellular Phylodynamic Analysis*** allows us to distill the features of the process. For example, should a part of the body come existence from ***z Founder Cells***, each of which is born at the same time, and has the same ***Cell-Heritable Cell Cycle Time, c***_***p***_, the part made of these ***z Founder Cells*** will also grow by the ***Cellular Allometric Growth Equation***. Even more complex examples of this type, including the process of ***ALLO-GROWTH*** in ***drosophila*** development, are amenable to ***Cellular Phylodynamic Analysis***, topics addressed in the APPENDIX.

#### Body part *ALLO-GROWTH* results from whole body *UNI-GROWTH*

Recall that the ***Cellular Allometric Growth Equation*** captures the rate of growth of the whole embryo, while the rate of growth for each body-part is:

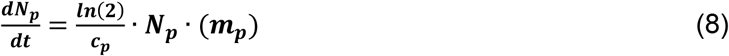

where ***c***_***p***_ is the ***Cell Cycle Time***, and ***m***_***p***_ the ***Mitotic Fraction*** of the cells of the body part. Thus, the ratio of the rate of growth of the body to the body-part is:

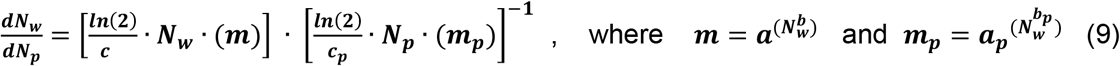

Consider the case where ***m***_***p***_***=m*** and ***c*** _***p***_***= c***, that is, where a body part has the same ***Mitotic Fraction*** and ***Cell Cycle Time*** as the embryo, and starts sometime later than the first cell:

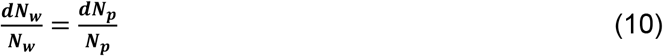

Integration reveals:

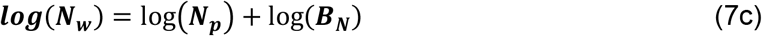

which is the allometric relationship, with ***S***_***N***_=**1**. Thus, growth of a part of the body by the ***Cellular Allometric Growth Equation*** is the consequence of ***Founder Cells*** giving rise to cell lineages in animals growing by the ***Universal Growth Equation***.

#### The *Cellular Allometric Slope, S_N_*, captures *Cell-Heritable* change in the *Cell Cycle Time, c_p_*, and body part size change by *Cellular-Selection*

Now, let us consider the case where ***m***_***p***_***=m*** but ***c*** _***p***_***≠ c***, that is, where a body part has the same ***Mitotic Fraction*** as the embryo but a different ***Cell Cycle Time***:

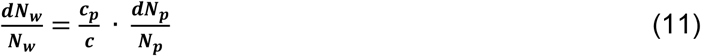

Integration reveals:

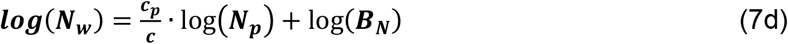

where:

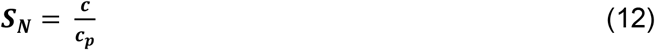

Thus, ***S***_***N***_, the ***Cellular Allometric Slope*** of the ***Cellular Allometric Growth Equation***, can be traced to the ***Cell Cycle Time, c***_***p***_ in each part of the body (FIGURE 3)!

On the other hand, similar analysis revealed the ***Mitotic Fraction, m***_***p***_, that is, a change in the fraction of cell dividing, to be an unlikely source for allometric log-linearity. Indeed, changes in the value of either ***a*** or ***b*** of the ***Universal Mitotic Fraction*** ***Equation*** (#4), which sets the value of ***m***_***p***_, does not result in straight lines on log-log graphs but in curves (not shown).

The changes in the value of ***c***_***p***_ seen by the ***Cellular Phylodynamic Analysis*** of body part growth are small (FIGURE 2 and FIGURES A8-A17 in APPENDIX). Thus, they have a minor impact on the average ***Cell Cycle Time, c***, in the embryo as a whole, but can have dramatic impacts on the relative sizes of body parts.

The value of ***S***_***N***_, the ***Cellular Allometric Slope*** of the ***Cellular Allometric Growth Equation***, appears on log-log graphs as a measure of whether a part of the body is growing faster (***S***_***N***_>**1**), slower (***S***_***N***_<**1**), or at the same speed (***S***_***N***_=**1**) as the body as a whole. Thus, ***ALLO-GROWTH***, and its mathematical abstraction, the ***Cellular Allometric Growth*** ***Equation*** (#7), distills ***Cellular-Selection***, that is, differential cellular proliferation,^15,16^ down to a single value, ***S***_***N***_. Embryos accomplish this by ***Cell-Heritable*** change in the ***Cell Cycle Time, c***_***p***._

What might be the mechanism of such a ***Cell-Heritable*** change in the ***Cell Cycle Time, c***_***p***_? DNA methylation has been found to induce ***Cell-Heritable*** changes in ***Cell Cycle Time***^48^, which one might imagine could occur at the ***Cellular Allometric Birth, B***_***N***_, of ***Founder Cells***, a hypothesis worthy of experimental examination.

#### CELLULAR POPULATION TREE VISUALIZATION SIMULATION

The reader will note that we have extracted information on the relationship between the number of cells in the body as a whole, ***N***_***w***_, with age, ***t***, from the cell lineage charts of nematodes (FIGURE 1). However, even without full lineage chart information, it is possible to reconstruct a plausible approximation of an animal’s cell lineage chart by a simulation which uses the growth equations of ***Cellular Phylodynamics*** and their parameters ***a, b, c, B***_***N***_, and ***S***_***N***_, extracted from batch growth data (for method, see APPENDIX).

In the FIGURE 4 (left) we display the idealized case of the cell lineage chart of an animal growing by the ***Universal Growth Equation***. Note how, as time goes on, an increasing fraction of cells, the ***Mitotic Fraction, m***, don’t divide, leading to some of the branches of the tree become extended in the absence of mitosis.

**FIGURE 4.**
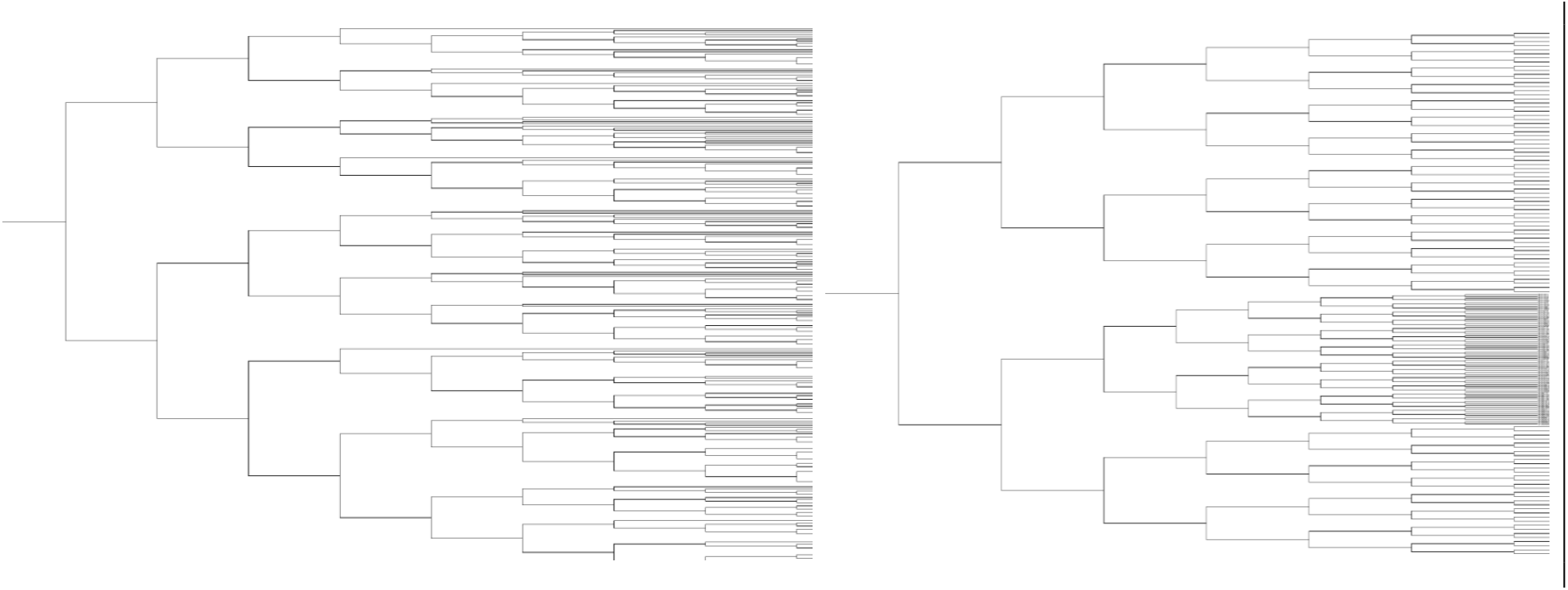
Cellular Population Tree Visualization Simulations. <u>Left,</u> The idealized case of the cell lineage chart of an animal growing by the ***Universal Growth Equation***. Note how, as time goes on, an increasing fraction of cells, the ***Mitotic Fraction, m***, don’t divide, and some of the branches of the tree become extended in the absence of mitosis. We have treated the ***Mitotic Fraction, m***, as a random variable, a plausible assumption simply for not knowing which cells are dividing at each moment in tim e, and a possible biological process, if the ***Mitotic Fraction*** is actually found to be the result of the discrete allocation of ligand molecule among cells, as indicated by the mathematical analysis of such a process (Equations #700-#720, APPENDIX). <u>Right,</u> The idealized case of the cell heritable shortening of the ***Cell Cycle Time*** that occurs in one ***Founder Cell*** when the embryo is 16 cells in size (***Cellular Allometric Birth, B***_***N***_=**8**); note how the progeny of the ***Founder Cell*** increased in number in comparison to the rest of the embryo. Code is executed in JAVA (see APPENDIX). Visualizations were made with IcyTree.^57^ Future generations of such simulations may well benefit for the stochastics simulator,^58^ located within BEAST.^59^

The ***Universal Growth*** and ***Cellular Allometric Growth Equations*** can also be taken to together, with their various parameters, to reconstruct an even more likely chart of an animal’s estimated cell lineage. In FIGURE 4 (right) we display the idealized case of the cell heritable shortening of the ***Cell Cycle Time*** that occurs in one ***Founder Cell*** when the embryo is 16 cells in size (***Cellular Allometric Birth, B***_***N***_=**16**); note how the progeny of the ***Founder Cell*** increased in number in comparison to the rest of the embryo. Left,

## Discussion

The ***Cellular Phylodynamic Analysis*** reported here, that is, the analysis of growth and development in units of numbers of cells, ***N***, has revealed two striking, and previously unappreciated aspects of embryonic life: ***UNI-GROWTH*** and ***ALLO-GROWTH***.

### * UNI-GROWTH

is the process by which growth slows as we increase in size. ***UNI-GROWTH*** is captured by the ***Universal Mitotic Fraction*** and ***Universal Growth Equations***, and their parameters, ***a, b***, and ***c***, the average ***Cell Cycle Time. UNI-GROWTH*** occurs by the decline in the fraction of cells dividing, the ***Mitotic Fraction***, occurring by the form of ***Universal Mitotic Fraction Equation, m=a^(N^b)***. From this expression, we could then derive, and test, the ***Universal Growth Equation***, whose accuracy was confirmed, from fertilization until maturity, for 13 species, including nematodes, mollusks, amphibians, fish, birds, rodents, cows, and humans.

### * ALLO-GROWTH

is the process by which body parts are created from ***Founder Cells. ALLO-GROWTH*** is captured by the ***Cellular Allometric*** and ***Allometric Growth Equations***, and their parameters, ***S, B, S***_***N***_, the ***Cellular Allometric Slope***, and ***B***_***N***_, the ***Cellular Allometric Birth. ALLO-GROWTH*** occurs by a ***Founder Cell*** acquiring a ***Cell-Heritable, Cell Cycle Time, c***_***p***_ at the ***Founder Cell’s Cellular Allometric Birth, B***_***N***_, and then undergoing mitotic expansion, as captured by the ***Cellular Allometric Slope, S***_***N***_, of the ***Cellular Allometric Growth Equation***. Whether a body part arises from a single ***Founder Cell***, or multiple ***Founder Cells*** arising at the same time with the same ***Cell Cycle Time***, growth by the ***Cellular Allometric*** and ***Allometric Growth Equations*** will occur. For body parts arising from a single ***Founder Cell***, the ***Cellular Allometric Birth, B***_***N***_, corresponds to the number of cells in the body when the ***Founder Cell*** arose.

For more than a century, biologists have appreciated that as we grow bigger, we also grow slower, that is, ***density dependent growth***, and that, the relative growth of parts of the body often form straight lines on log-log graphs, that is, ***allometric growth***.^3,6,7,13^ ***Cellular Phylodynamic Analysis*** captured these two observations in ***UNI-GROWTH*** and ***ALLO-GROWTH***, and uncovered the surprising finding that they are really different manifestations of the same process, since the ***Cellular Allometric Growth Equation*** can be derived from the ***Universal Growth Equation***.

***UNI-GROWTH*** and ***ALLO-GROWTH***, capture, in mathematical form, ***Cellular-Selection***, that is, differential cellular proliferation.^15,16^ The ***Cellular Allometric Slope, S***_***N***_, of the ***Cellular Allometric Growth Equation*** captures ***Cell-Selection*** in the increase or decrease in the size of a body part, and traces its origin to the ***Cell Cycle Time, c***_***p***_. The ***Mitotic Fraction, m***, of the ***Universal Mitotic Fraction Equation*** captures ***Cell-Selection*** in the restriction of cell division to a selected few cells. The ***Cellular Allometric Birth, B***_***N***_, of the ***Cellular Allometric Growth Equation*** captures ***Cell-Selection*** in the choice of ***Founder Cells*** among all the other cells of the embryo. Thus, these ***Cellular Phylodynamic*** equations, and their parameters, provide quantitative metrics for ***Cellular-Selection*** in its role in the creation of anatomical structures by outgrowth from the undifferentiated body mass, and in the molding of the fine scale detail of anatomy, processes that lie at the heart of embryonic development.^15,16^

***ALLO-GROWTH*** also captures, in mathematical form, ***Cell-Heritability***, the persistence of phenotypic characteristics after cell division.^15,16^ The ***Cellular Allometric Slope, S***_***N***_, of the ***Cellular Allometric Growth Equation*** captures the ***Cell-Heritable*** change in the ***Cell Cycle Time, c***_***p***_, that occurs at the ***Cellular Allometric Birth, B***_***N***_, of the ***Founder Cell***. However, the ***Cellular Phylodynamic Analysis*** of growth and development in units of numbers of cells, ***N***, that we describe here has not yet told us how ***Founder Cells*** acquire such a ***Cell-Heritable*** change. Of course, other cellular processes also display ***Cell-Heritability***, such as gene expression, cell size, and the geometry of each cell division, and how such programs of mitotic geometry contribute to the shape of the animal.^49^ Details for how the ***Cellular Phylodynamic Analysis*** of such processes could be carried out computationally are presented in the APPENDIX, where we outline a technique, the ***BinaryCellName*** method, for analysis of cell lineages at the single cell level, and another technique, the ***S***_***N3***_ ***Method***, for analysis of the cellular basis of shape formation at the macroscopic level.

Growth is critical to the survival and evolution of individuals and species. Human fetuses that grow to average birthweight in 8 months, or 10 months, suffer a heart-breaking increase in its risk of disability and death. In the APPENDIX, we show how ***Cellular Phylodynamic*** mathematics can provide equations for the assessment and management of human fetal size and growth. ***Life History Theory*** teaches that in nature the struggle for existence includes growing faster or slower or bigger or smaller than competitors, causing animals to acquire growth curves molded by the cruel disciple of Darwinism.^29,30^ The ***Universal Growth Equation***, being linked to the developmental biology and genetics of the creatures that must adapt, gives us a tool for achieving ever greater biological reality in our understanding of the impact of individual growth on populations.

Growth and development are not only crown fascinations of embryology, they also present practical challenges in agriculture and aquaculture. In the APPENDIX, we outline how the ***Cellular Phylodynamic Analysis*** can be put to work in breeding fish with optimal growth characteristics for use in the aquaculture industry.

We have seen that ***Cellular Phylodynamic Analysis*** can be carried out with cell numbers, ***N***_***p***_ and ***N***_***w***_, counted from in cell lineage charts. From the number of these cells, we could identify the form and parameters of the ***Universal Mitotic Fraction, Universal Growth***, and ***Cellular Allometric Growth Equations***. We have also seen that we can reverse this line of reasoning, so that even in the absence of such lineage charts, useful approximations of cell lineage can be constructed by simulation, utilizing the parameters of these equations extracted from batch growth data. Following the thinking of Stadler, Pybus, and Stumpf,^22^ we call these “***Cellular Population Tree Visualization Simulations***”, which should bring the powerful mathematics of lineage formation, coalescent theory,^22–27^ to developmental biology. For example, there has been much interest in the past few years in extracting information on gene expression, differentiation, and lineage formation, from single cell RNA expression,^50,53,54^ cellular barcoding,^21,51^ and somatic mutation.^52^ However, as Telford^21^ and Klein^53,54^ have pointed out, without knowledge of the mitotic pattern of an animal’s cell lineage, extraction of useful information from such single cell data is problematic. One might hope, and indeed expect, that the potential of ***Cellular Population Tree Visualization Simulation***, driven by the parameters measured by ***Cellular Phylodynamic Analysis***, could unburden single cell analysis from the requirement of deciphering ancestry, and unlock the much hoped-for treasure of how gene expression is linked to cell lineage formation.

The ability of cells to slow down mitosis, so as to glide to ideal body size, and gear up mitosis, so as to create useful parts, are essential qualities of animal life. By the simple examination in units of numbers of cells, ***N***, we have seen these qualities in ***UNI-GROWTH*** and ***ALLO-GROWTH***, and their defining abstractions, the ***Universal Mitotic Fraction, Universal Growth***, and ***Cellular Allometric Growth Equations***, in animals as diverse as tunicates, vertebrates, mollusks, and nematodes. The forms of these equations provide hints of their biochemical mechanisms, and their broad phylogeny suggest invention at the dawn of animal life.^55,56^ If so, perhaps ***Cellular Phylodynamic Analysis*** is giving us a peek at the mysterious billion-year-old spectacle of how our cells become our selves.

**TABLE.**
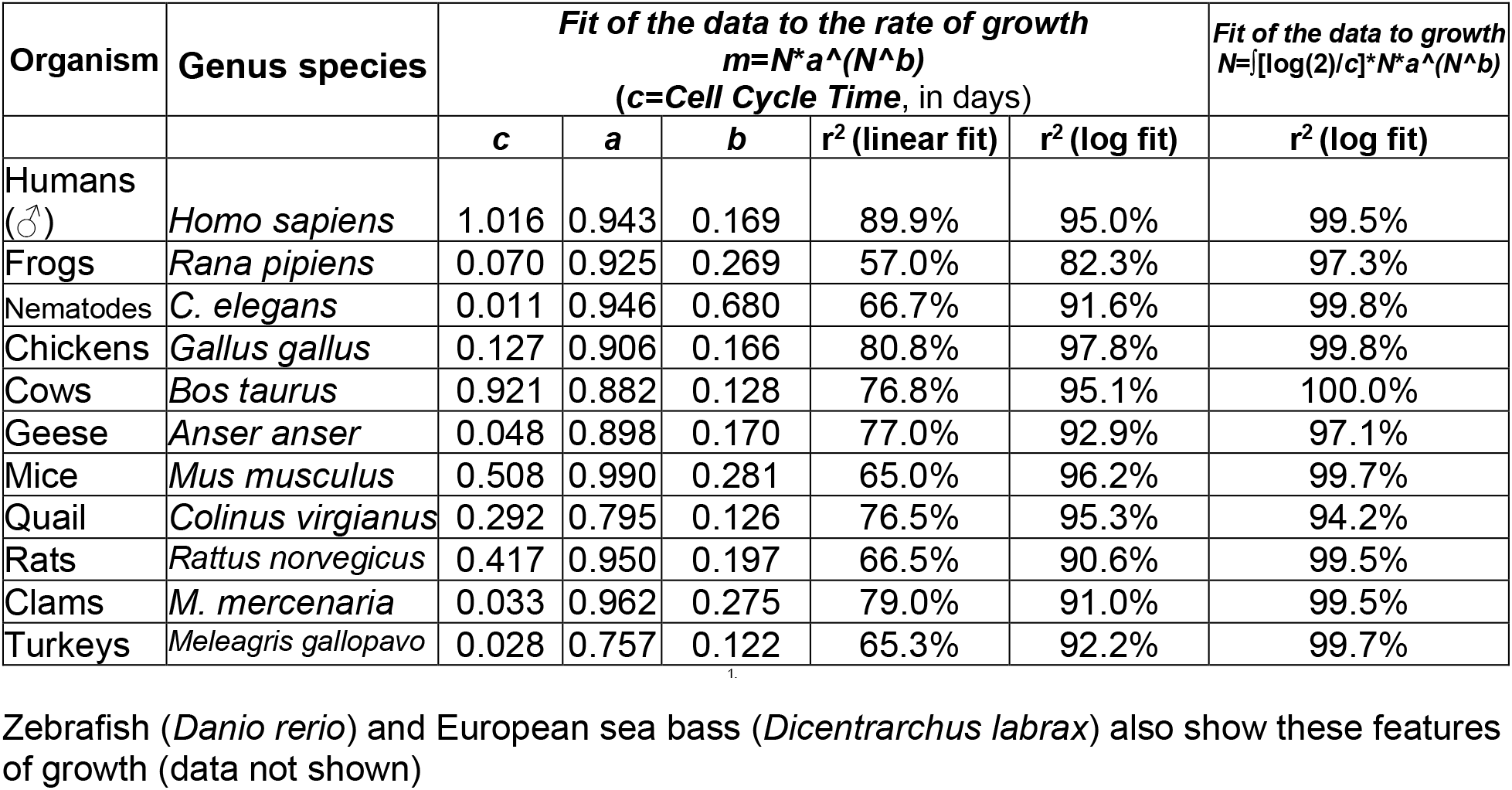
Parameters and goodness-of-fit metrics to the ***Universal Growth*** and ***Universal Mitotic Fraction Equations***.

## Acknowledgments

We thank Alain Goriely, Derek Moulton, Aris Papageorghiou, Jose Villar, Stephen Kennedy, Eleonora Staines Urias, Henning Tiemeier, and Hari Shroff for many useful suggestions and discussions, and Alex Gould for kindly sharing data.

